# Honokiol decreases alpha-synuclein mRNA levels and reveals novel targets for modulating alpha-synuclein expression

**DOI:** 10.1101/2023.07.06.547988

**Authors:** Sara Fagen, Jeremy D. Burgess, Melina Lim, Danilyn Amerna, Zeynep B. Kaya, Ayman H. Faroqi, Priyanka Perisetla, Natasha N. DeMeo, Iva Stojkovska, Drew J. Quiriconi, Joseph R. Mazzulli, Marion Delenclos, Suelen L. Boschen, Pamela J. McLean

**Affiliations:** Department of Neuroscience, Mayo Clinic, Jacksonville, FL, USA; Mayo Clinic Graduate School of Biomedical Sciences, Mayo Clinic College of Medicine, Rochester, MN, USA; Feinberg School of Medicine, Northwestern University, Chicago, IL, USA; Department of Neurosurgery, Mayo Clinic, Jacksonville, FL, USA

**Keywords:** Alpha-synuclein (αSyn), Parkinson’s disease, *SNCA*, natural compound, polyphenol, therapeutic target

## Abstract

Neuronal inclusions comprised of aggregated alpha-synuclein (αsyn) represent a key histopathological feature of neurological disorders collectively termed “synucleinopathies”, which includes Parkinson’s disease (PD). Mutations and amplifications in the *SNCA* gene encoding αsyn cause familial forms of PD and a large body of evidence indicate a correlation between αsyn accumulation and disease. Decreasing αsyn expression is recognized as a valid target for PD therapeutics, with down-regulation of *SNCA* expression potentially attenuating downstream cascades of pathologic events. Honokiol (HKL) is a polyphenolic compound derived from magnolia tree bark that has demonstrated neuroprotective properties. Here, we describe potential beneficial effects of HKL on αsyn levels in multiple experimental models. Using human neuroglioma cells stably overexpressing αsyn and mouse primary neurons, we demonstrate that HKL treatment results in a significant decrease in αsyn expression at both the protein and mRNA levels. Our data support a mechanism whereby HKL acts by post-transcriptional modulation of *SNCA* rather than modulating αsyn protein degradation. Additionally, transcriptional profiling of mouse cortical neurons treated with HKL identified several differentially expressed genes (DEG) as potential targets to modulate *SNCA* expression. Overall, these data highlight a viable strategy to reduce αsyn levels, which represents a promising target to modify disease progression in PD and other synucleinopathies. In addition, HKL acts as a powerful tool for investigating *SNCA* gene modulation and its downstream effects.

## 1. Introduction

Alpha-synuclein (αsyn) accumulation is a key feature in the pathogenesis of Parkinson’s disease (PD) and related synucleinopathies (Spillantini, Schmidt et al. 1997). These diseases are characterized by the misfolding and aggregation of αsyn protein that can propagate between cells in the brain and accumulate as Lewy bodies (LB) and Lewy neurites (LN) in susceptible cellular populations (Luk, Kehm et al. 2012). Expression of αsyn is a strong disease modifier as individuals with a triplication of the *SNCA* gene locus develop aggressive forms of PD with dementia (Polymeropoulos, Lavedan et al. 1997, Singleton, Farrer et al. 2003, Houlden and Singleton 2012). Experimentally, the expression level of αsyn is an important determinant of the rate of fibrillization and neurotoxicity (Rockenstein, Nuber et al. 2014). Additionally, knocking out αsyn in mice (Dauer, Kholodilov et al. 2002) or knocking down αsyn in differentiated human dopaminergic cells increases resistance to the mitochondrial toxin MPP+ (Fountaine and Wade-Martins 2007). Disease-modifying therapies for PD remain a major unmet medical need and reducing αsyn levels is a promising therapeutic target (Junn, Lee et al. 2009, Mandler, Valera et al. 2015, Schneeberger, Tierney et al. 2016, Valera, Spencer et al. 2016, Kallab, Herrera-Vaquero et al. 2018). Downregulation of αsyn via the use of passive or active immunization (Helmschrodt, Hobel et al. 2017, Kantor, Tagliafierro et al. 2018, Brys, Fanning et al. 2019, Savitt and Jankovic 2019, Uehara, Choong et al. 2019), antisense oligonucleotides strategy (Junn, Lee et al. 2009, Dehay, Vila et al. 2016, Vaikath, Hmila et al. 2019), and viral vector technology (Menon, Kofoed et al. 2021) has demonstrated beneficial effects and could attenuate downstream cascades of pathologic events. However, neurotoxicity associated with robust reduction of *SNCA* mRNA levels was reported in studies that utilized RNAi tools to directly target *SNCA* transcripts and immunization against αsyn still requires larger efficacy trials. Therefore, further studies are necessary to develop a strategy that safely and successfully down-regulates αsyn.

Phytochemical compounds can contribute to the brain’s chemical balance and current evidence supports the applicability of natural compounds to treat neurodegenerative disorders (Perez-Hernandez, Zaldivar-Machorro et al. 2016, Sharifi-Rad, Lankatillake et al. 2020). The medicinal properties of plants are mostly attributed to their secondary phytochemical metabolites that have a wide spectrum of pharmacological activities, including but not limited to, antioxidant, anti-tumor, anti-inflammatory, and neuroprotective properties (Kumar and Khanum 2012, Forni, Facchiano et al. 2019). Honokiol (HKL) is a polyphenolic compound derived from the bark of magnolia plant that has demonstrated favorable effects in experimental models of cancer (Ong, Lee et al. 2019), Alzheimer’s disease (Wang, Dong et al. 2018) (Ramesh, Govindarajulu et al. 2018), and PD (Chen, Chang et al. 2018). Additionally, oral administration of HKL attenuated age-related cognitive impairment and neuronal injury in senescence accelerated mice (Matsui, Takahashi et al. 2009). Importantly, intraperitoneal administration of HKL produced a desirable bioavailability (Wang, Duan et al. 2011) with considerable blood brain barrier penetration (Lin, Chen et al. 2012).

A previous study demonstrated that chronic HKL treatment prevented dopaminergic neuronal loss and motor impairments in a hemi-parkinsonian mouse model (Chen, Chang et al. 2018). Here, we use multiple cellular models to further characterize potential benefits of HKL in PD. Additionally, we demonstrate that HKL can modulate *SNCA* expression levels and interrogate the molecular mechanism(s) whereby HKL has its effect on *SNCA*.

## 2. Materials and methods

### 2.1. Wt-αsyn H4 cell culture

A stable H4 neuroglioma cell line expressing human wt-αsyn was generated and previously described (Moussaud, Malany et al. 2015). Cells were maintained at 37°C in a 95% air/5% CO2 humidified incubator in Opti-MEM supplemented with 10% FBS, 200µg/ml G418, and 200µg/ml Hygromycin. Tetracycline (1µg/ml, Sigma, #T7660-5G) was added to culture media to block the expression of αsyn in the transgene cells.

### 2.2. Cortical primary neurons preparation

Pregnant adult CD-1 mice were ordered from Jackson Laboratory (Bar Harbor, ME). Cell culture dishes were freshly prepared for each litter and were coated with poly-D-lysine (PDL) diluted in DPBS at a final concentration of 0.1 mg/ml. All media were made fresh for each litter and used within 1-2 weeks. Dissection buffer was prepared containing 1X HBSS without phenol red, calcium, or magnesium (Gibco, #14185-052) and HEPES (Gibco, #15630-106) at final concentration of 10mM. Primary culture FBS medium comprised NeuroBasal Medium without L-Glutamine (Gibco, #21103-049), 10% FBS (Gibco, #10437-028), 1% GlutaMAX (Gibco, #35050-061), and 1% Penstrep (Gibco, #15140-122). Neuronal media comprised NeuroBasal medium without L-Glutamine, 2% B-27 (ThermoFisher, # 17504044), 1% GlutaMAX, and 1% Penstrep.

### 2.3. HKL and DMSO solutions preparation

A 100mM stock solution of HKL (MedChem express, #HY-N0003-50MG,) was dissolved in 100% DMSO and stored at −30^ο^C. For treatment of cells, HKL was further diluted in the appropriate cell culture media at a final concentration of 10µM. DMSO controls were similarly prepared with a final concentration of 0.01% DMSO. Wt-αsyn and primary cortical neurons were treated with 0.01% DMSO or 10µM HKL for 72 hours before being processed.

### 2.4. Induced pluripotent stem cell (IPSC)-derived neurons treatment with HKL

Lymphoblast cell lines were acquired from the Coriell Institute for Medical Research, Line # GM15010, female origin (New Jersey, USA) from a patient carrying a triplication in the *SNCA* gene, reprogrammed into induced pluripotent stem cells (iPSCs) called line ‘3x-1’, and characterized previously (Stojkovska *et al*., 2022). iPSCs were cultured on Matrigel (Corning, #354277) coated plates and maintained in mTESR1 media. Differentiation into midbrain dopaminergic neurons occurred using previously established protocols (Kriks et al, Nature, 2011). Briefly, iPSC lines were accutased (Corning, # 25058CI) and seeded onto Matrigel (Corning, #354277) coated plates, allowed to grow to confluency, then treated with dual SMAD inhibitors followed by a cocktail of growth factors (Cuddy et al, Neuron 2019). After differentiation, the neurons were cultured in neurobasal medium (ThermoFisher, # 21103049) with NeuroCult™ SM1 Neuronal Supplement (Stem cell technologies, #5711) and 1% glutamine and penicillin / streptomycin. Patient derived *SNCA* triplication IPSC-neurons were used to evaluate the effects of HKL in a human relevant model of synucleinopathy. Neurons were matured for 60 days and subsequently treated with 10µM HKL or 0.01% DMSO for 72 hours. Cells were pelleted, snap frozen, and shipped to Mayo Clinic-Jacksonville for αsyn and *SNCA* mRNA quantitation.

### 2.5. Western blotting analysis

To prepare whole cell lysates, cells were washed twice with ice-cold PBS and total proteins were isolated by incubating the cells on ice in radio-immunoprecipitation assay (RIPA) lysis buffer (50mM Tris–HCl, pH 7.4, 150mM NaCl, 1mM EDTA, 1mM EGTA, 1.2% Triton X-100, 0.5% sodium deoxycholate, and 0.1% SDS) containing 1 mM phenylmethylsulfonyl fluoride (PMSF), protease inhibitor cocktail, and phosphatase inhibitor cocktail. Collected cells were sonicated on ice and centrifuged at 10,000 × g for 10min at 4°C. The protein concentration was determined with Bradford reagent (Thermo Fisher, #23225). 5-10μg of total proteins were separated on bis-tris polyacrylamide gradient gels (NuPAGE Novex 4-12% Bis-Tris Gel, Life tech, #NW04120BOX) or TGX stain-free gels (BioRad, #4568126) and transferred to PVDF membranes. Membranes were then blocked for 1h at room temperature (RT) in TBS-T (500mM NaCl, 20mM Tris, 0.1% Tween 20, pH 7.4) supplemented with 5% non-fat dried milk. Subsequently membranes were incubated overnight at 4°C with primary antibodies (see Table 1 for antibody list) followed by 1h at RT with HRP-conjugated secondary antibodies. Proteins were detected using an enhanced chemiluminescent detection system (ECL, EMD Millipore, #WBKLS0500) and the BioRad ChemiDoc MP (#12003153) imaging system. Blots were quantified using ImageJ and Image lab software (BioRad, #17006130) and normalized to the appropriate loading control such as Vinculin (Sigma, #V9131), GAPDH (Abgent, #AP7873a), Actin (Sigma, #A5060), or total protein.

### 2.6. Cell toxicity, viability, and proliferation

Cell toxicity was assessed using the Toxilight bioassay kit (Lonza, #LT17-217) in both primary cortical neurons and in wt-αsyn cells to determine the viability of cells after treatment with HKL. In both cases, cells were treated with 0.01% DMSO or 10uM HKL for 72hrs. The culture plates were removed from the 37°C incubator and left at room temperature for 5 minutes. 20µl of conditioned media was transferred from each well to a 96-well luminescence compatible plate. Fresh adenylate kinase (AK) detection reagent was used for each experiment, in which 100µl was added to each conditioned media containing well and incubated at room temperature for 5 minutes. Luminescence was then read for 1 second in a microplate reader (EnVision, PerkinElmer).

Cell proliferation was determined using WST-1 (abcam, #ab155902) assay according to manufacturer instructions. Briefly, WT-asyn cells were plated in a 96-well plate with 10,000 cells per well in 100µl of media containing tetracycline. The following day, tetracycline was removed, and the cells were treated with either 0.01% DMSO or 10µM HKL in 100µl of media. After 24, 48, and 72 hours, 10 µl/well of WST-1 reagent was added to each well and the plate was incubated at 37°C for 4 hours. WST-1 absorbance was read at 450nm on an EnVision microplate reader (EnVision, PerkinElmer). Three biological replicates of the cell proliferation assay were performed each with three technical replicates for each group in each set.

### 2.7. Degradation assays

Wt-αsyn cells were grown in the absence of tetracycline for 24 hours. Then, tetracycline was added to suppress further expression of αsyn, and cells were treated with 10µM HKL or 0.01% DMSO. Cells were then harvested at 0, 6, 12, 24, 48, and 72 hours for western blot and qPCR analysis to evaluate rate of αsyn and *SNCA* mRNA degradation.

### 2.8. mRNA extraction and qPCR

Total RNA was extracted from cells using TRIzol Reagent (Ambion Life Technology, #15596018) followed by DNase RNA cleanup using RNeasy (Qiagen, #74106). The quantity and quality of RNA samples were determined by the Agilent 2100 Bioanalyzer using an Agilent RNA 6000 Nano Chip. Complementary DNA (cDNA) synthesized with Applied Biosystems High-Capacity cDNA Archive Kit was used as a template for relative quantitative PCR using ABI TaqMan chemistry (Applied Biosystems, #4368814). mRNA expression was quantified using Hs00240906_m1 (human *SNCA)*, Mm01188700_m1(mouse *Snca*), Mm00497442 _m1 (*txnl1,* Thioredoxin-Related Protein 1), and Hs02800695_m1(*HPRT1*, Hypoxanthine-guanine phosphoribosyltransferase) probes. Each sample was run in quadruplet replicates on the QuantStudio 7 Real-Time PCR System (Thermo Fisher) and quantification was done using the 2–ΔΔCT method.

### 2.9. RNAscope/immunocytochemistry (ICC)

RNAscope is a variation of FISH (fluorescent *in situ* hybridization) used to visualize RNA transcripts within cells. Kits and probes were purchased from ACD (Fluorescent multiplex detection reagent kit, #320851 and *SNCA*, #571241) and used according to manufacturer’s instructions. Briefly, cells were fixed with 4% PFA, *SNCA* mRNA was amplified, and the secondary fluorescent detection probe (Thermo Fisher, Alexa Fluor 488 #A11000) was added. Cells then underwent ICC to examine αsyn protein levels. Cells were permeabilized using 0.1% Triton-X in 1X PBS and incubated at RT. Following additional washes, bovine serum albumin (BSA) was used to block non-specific antigens. Primary antibody, 4B12 (αsyn, # 807802, 1:1,000) was diluted in blocking buffer and incubated at RT. Secondary fluorescent antibody Alexa Fluor 568 (Thermo Fisher, # A11004, 1:1,000), were incubated after washes in the dark. Hoechst staining (#H3570, 1:5,000) was completed prior to imaging and quantification on a Perkin Elmer Operetta CLS High Content Imager (Johns Creek, GA).

### 2.10. Honokiol derivatives

Magnolol (#M0125) and 4-0-Methylhonokiol (#M184770) were ordered from LKT Labs (St. Paul, MN). Derivatives arrived in powered form and were prepared in the same manner as HKL (i.e., dissolved in 100% DMSO and diluted to 10µM in the appropriate culture medium).

### 2.11. RNA sequencing

The mRNA samples were sequenced by the Mayo Clinic Genome Analysis Core (Rochester, MN) using Illumina HiSeq 4000 (San Diego, CA). Reads were mapped to the mouse genome mm10. The raw gene read counts, along with sequencing quality control, were generated using the Mayo Clinic RNA sequencing (RNA-seq) analytic pipeline: MAP-RSeq version 3.0.1. Conditional quantile normalization (CQN) was performed on raw gene counts to remove biases created by GC content and technical variation, to adjust for gene length and library size differences, and to obtain similar quantile-by-quantile distributions of gene expression across samples. Based on the bimodal distribution of the CQN-normalized and log2-transformed reads per kilobase per million (RPKM) gene expression values, genes with average log2 RPKM ≥2 in at least one group were considered to have expression above the detection threshold. Using this selection threshold, 19,005 genes were included in downstream analyses.

### 2.12. Rotenone treatment

Wt-αsyn cells were plated in 6-well plates at 1×10^5^ cells/well with tetracycline. The following day, tetracycline was removed and 0.5μM rotenone (#R8875,Sigma) was added for 2 hours at 37°C. After the 2-hour incubation, cells were washed, fresh media was added, and wells were treated with 0.01% DMSO or 10μM HKL for 72 hours then harvested for protein and mRNA analysis.

### 2.13. Statistical analysis

All data were assessed using Graph Pad Prism 9 software (San Diego, CA) and analyzed by one-way ANOVA with Dunnett’s multiple comparisons test or unpaired *Student t* test where appropriate. Proliferation rate of wt-αsyn cells was analyzed with repeated measures two-way ANOVA, followed by Dunnett’s multiple comparisons test. Statistical analysis of degradation rate of αsyn and *SNCA* mRNA was conducted with paired *Student t* test. Differences were considered statistically significant when *p* < 0.05. Results are presented as mean ± standard error of the mean (SEM).

## 3. Results

### 3.1. HKL reduces overexpressed αsyn *in vitro*

Because HKL has previously been reported to inhibit αsyn amyloid fibril formation in a cell-free aggregation assay (Das, Pukala et al. 2018) we sought to examine the effect of HKL on αsyn in cellular synucleinopathy models given the critical role this protein plays in PD pathogenesis. Tetracycline-regulated H4 neuroglioma cells stably overexpressing human wt-αsyn were treated with escalating doses of HKL for 72 hours, harvested, and αsyn protein levels were assessed by Western blot. While doses of HKL in the range 0.625 μM – 5 μM did not alter αsyn levels, 10 μM HKL decreased αsyn protein levels by 39% (F (7,16) = 21.84, p < 0.0001) (Figure 1A). Therefore, we chose to use 10μM HKL for the remainder of the experiments. Additional experiments confirmed that 10μM HKL consistently reduces αsyn levels by up to 70% (*t* (14) = 9.18, p < 0.0001) (Figure 1B). Because αsyn expression in the H4 stable cell line is under control of a constitutively active tetracycline regulated promoter, in support of a specific effect on αsyn expression, we confirmed that HKL has no effect on expression of GFP in a similar tetracycline-regulated GFP-expressing H4 stable cell line (Supplemental Figure S1). Importantly, 10µM HKL treatment did not induce toxicity and did not affect cell proliferation in H4 wt-αsyn overexpressing cells (Supplemental Figure S2A-C). Together, these findings support HKL as an effective and safe compound to modulate αsyn levels *in vitro*.

**Figure 1.**
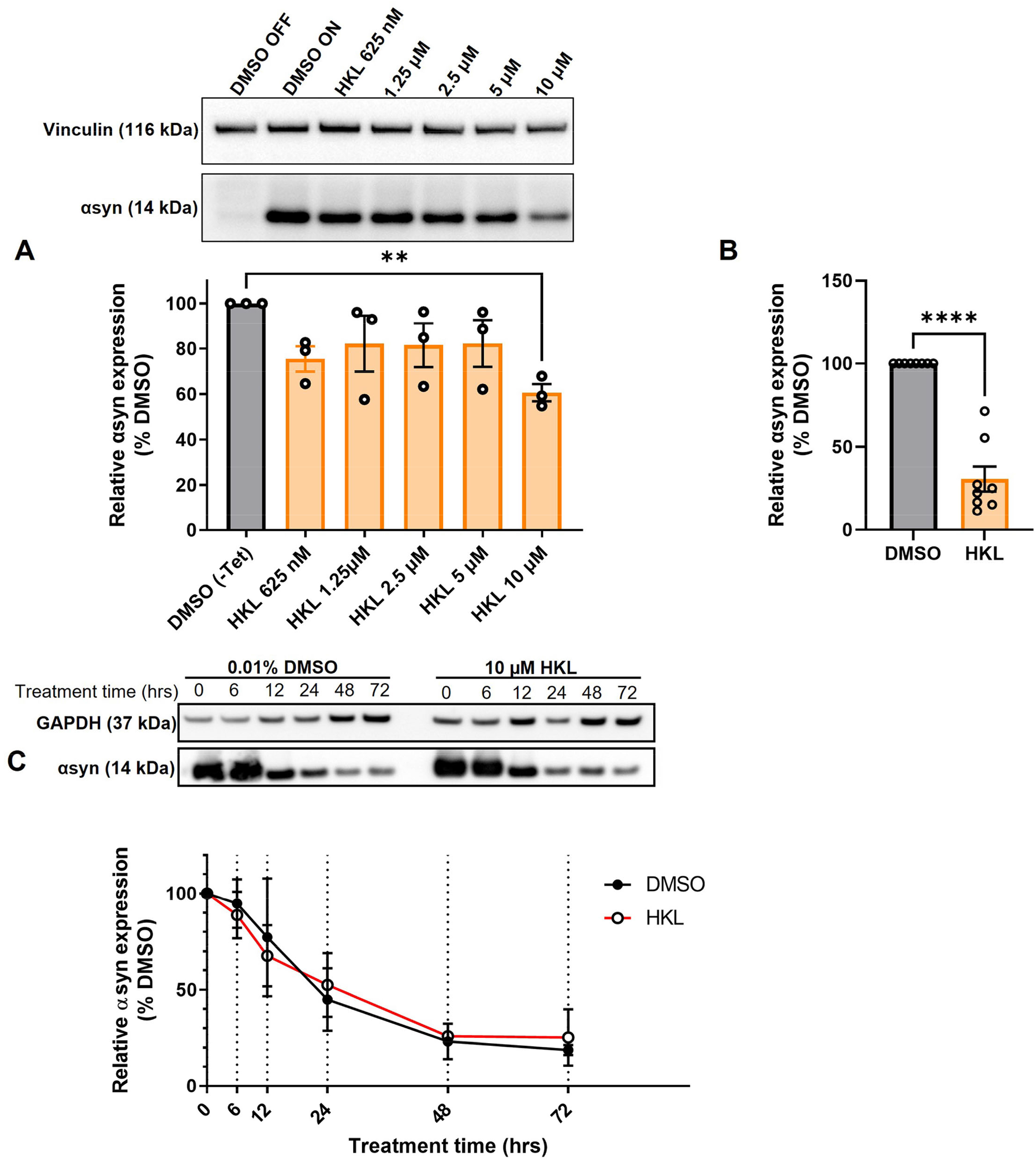
HKL reduces αsyn levels in H4 cells stably overexpressing wt-αsyn. Western blot and quantification of αsyn expression following treatment with different doses of HKL (n = 3 biological replicates/treatment, One-way ANOVA and Dunnet’s post-hoc) (A). Confirmation of αsyn expression reduction induced by 10µM HKL (n = 8 biological replicates/treatment, unpaired Student’s *t* test) (B). Western blot and quantification of αsyn levels at 0, 6, 12, 24, 48, and 72h of 10µM HKL treatment (n = 3 biological replicates/time point, paired Student’s *t* test) (C). Data are represented as mean ± SEM. **p < 0.01; ****p < 0.0001.

### 3.2. HKL does not increase αsyn degradation

One mechanism by which αsyn expression may be decreased by HKL is via an increase in the rate of protein degradation. Next, we conducted a degradation assay in wt-αsyn cells to determine if increased degradation was the primary mechanism by which HKL regulated αsyn levels (Figure 1C). Here we took advantage of tetracycline regulation to study the degradation rate of αsyn in the presence of HKL or vehicle (DMSO). Cells overexpressing wt-αsyn were cultured in the absence of tetracycline to allow αsyn expression before being treated with 10µM HKL. At time zero, tetracycline was added to the media to inhibit additional gene expression and samples were collected at 0, 6,12, 24, 48, and 72 hours post treatment. Interestingly, HKL did not change the degradation rate of αsyn (t (5) = 0.068, p = 0.95) compared to vehicle treated cells.

### 3.3. HKL reduces *SNCA* transcription in cells overexpressing wt-αsyn

If decreased protein expression in the presence of HKL is not due to an increase in rate of degradation, an alternative explanation could be that regulation is occurring at the level of transcription. To test this, we treated H4 cells overexpressing wt-αsyn with HKL for 72 hours and assessed *SNCA* mRNA levels using quantitative PCR. We observed a significant decrease (51%) in *SNCA* transcripts levels in cells treated with HKL compared to vehicle (*t* (14) =5.35, p < 0.0001) (Figure 2A). This decrease was confirmed using RNAscope to visualize and quantify *SNCA* transcripts and multiplexed with ICC to evaluate corresponding αsyn protein (Figure 2B). We calculated the total number of nuclei per field (n = 25 fields) and normalized the average *SNCA* mRNA spots and the average αsyn protein fluorescence intensity to the average number of nuclei for each treatment. Taken together, qPCR, RNAscope, and ICC confirm that HKL treatment significantly decreases *SNCA* transcript levels by 73% (*t* (6) = 18.85, p < 0.0001) and αsyn expression by 45% (*t* (6) = 3.94, p < 0.01) while not affecting expression of αTubulin (*t* (6) = 2.20, p = 0.07) (Figures 2C-E).

**Figure 2.**
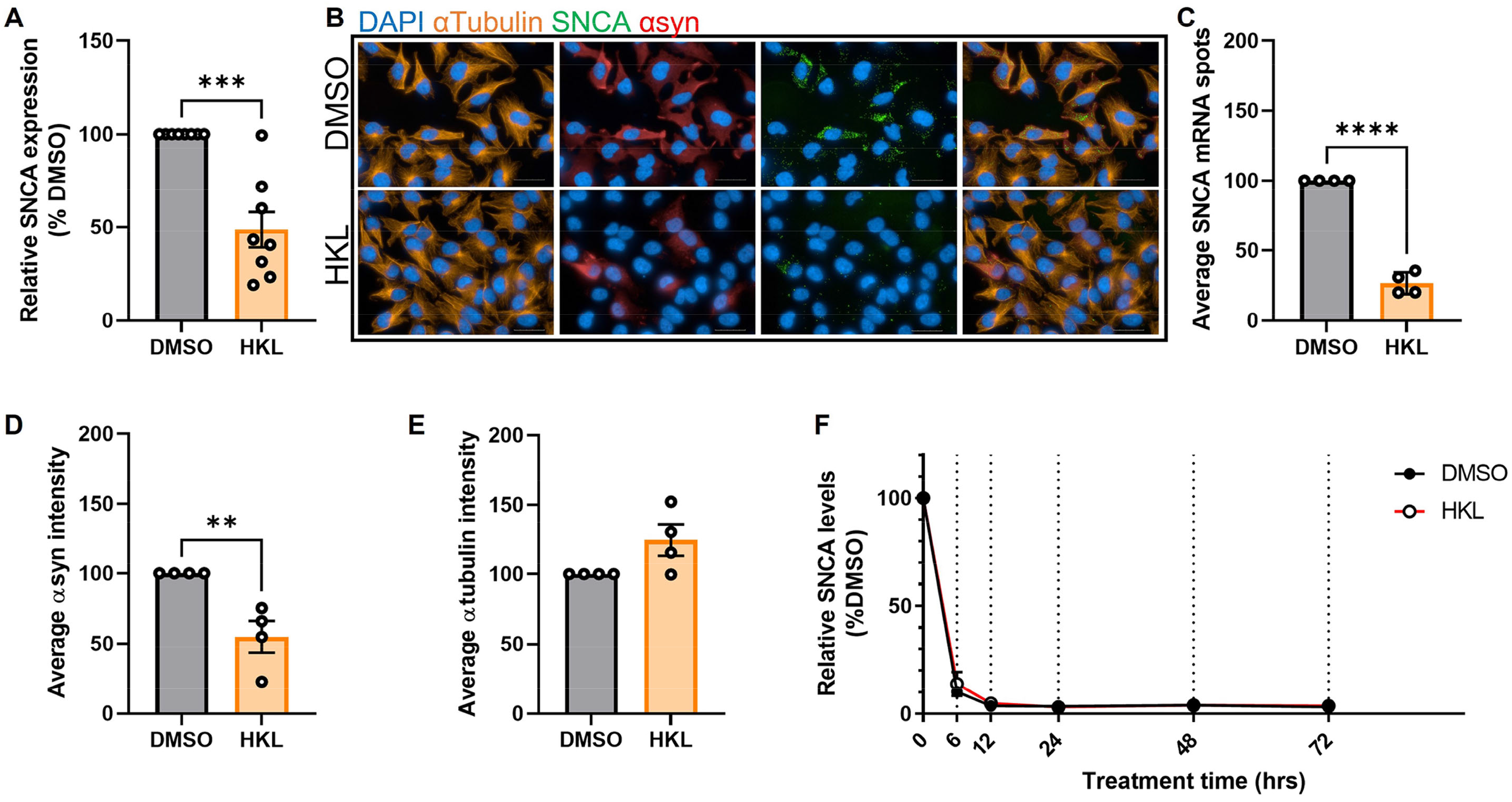
HKL reduces expression of αsyn transcripts. qPCR assay to determine *SNCA* mRNA levels after 10µM HKL treatment of H4 wt-αsyn cells (n = 8 biological replicates/treatment) (A). Representative RNAscope images of H4 wt-αsyn cells stained for α-tubulin, *SNCA* mRNA, and αsyn (B). Four biological replicates with 25 fields/well (2 wells/replicate) were evaluated to determine the effects of HKL and vehicle on *SNCA* mRNA (C), αsyn (D), and α-tubulin (E). Quantification of *SNCA* mRNA levels at 0, 6, 12, 24, 48, and 72h of 10µM HKL treatment in wt-αsyn cells (n = 3 biological replicates/time point, paired Student’s *t* test) (F). Data are analyzed with unpaired Student’s *t* test and are represented as mean ± SEM. **p < 0.01, ***p < 0.001, ****p < 0.0001. Scale bars = 200 µm

### 3.4. HKL does not affect rate of *SNCA* mRNA degradation

Our data so far indicate that HKL can modulate αsyn expression and that the modulation is via a transcription related mechanism. The nature of the H4 overexpressing cells (wt-αsyn) excludes the possibility that regulation is at the level of the promoter; thus, we examined whether HKL modulates *SNCA* levels post-transcriptionally. Here we again took advantage of our tetracycline regulated cell lines to evaluate the rate of mRNA degradation using the same paradigm used previously to examine rate of protein degradation. Interestingly, we determined that HKL does not affect the degradation rate of *SNCA* mRNA compared to vehicle control (Figure 2F) (*t* (5) = 1.38, p = 0.22).

### 3.5. HKL reduces αsyn in mouse primary cortical and but not in iPSC-derived neurons

Because our stable cell lines expressing wt-αsyn are under the control of a constitutively active promoter yet our data support HKL reducing expression by altering levels of transcription, we examined the effect of HKL on endogenous levels of αsyn where expression is controlled by the endogenous promoter. To evaluate the effect of HKL on endogenous αsyn expression, mouse primary cortical neurons were treated at 7 days-in-vitro with escalating doses of HKL for 72 hours. Interestingly, HKL reduced *Snca* mRNA levels in mouse primary cortical neurons at a dose as low as 6µM (Figure 3A) (F (5,12) = 8.67, p < 0.01). To be consistent, however, we continued to use 10µM HKL in subsequent experiments. Consistent with our previous data, 10μM HKL treatment resulted in a 44% decrease in αsyn protein expression (Figure 3B) (*t* (10) = 9.05, p < 0.0001) and a 25% decrease in *Snca* mRNA levels (Figure 3C), (*t* (16) = 7.16, p < 0.0001) compared to vehicle (DMSO) treatment.

**Figure 3.**
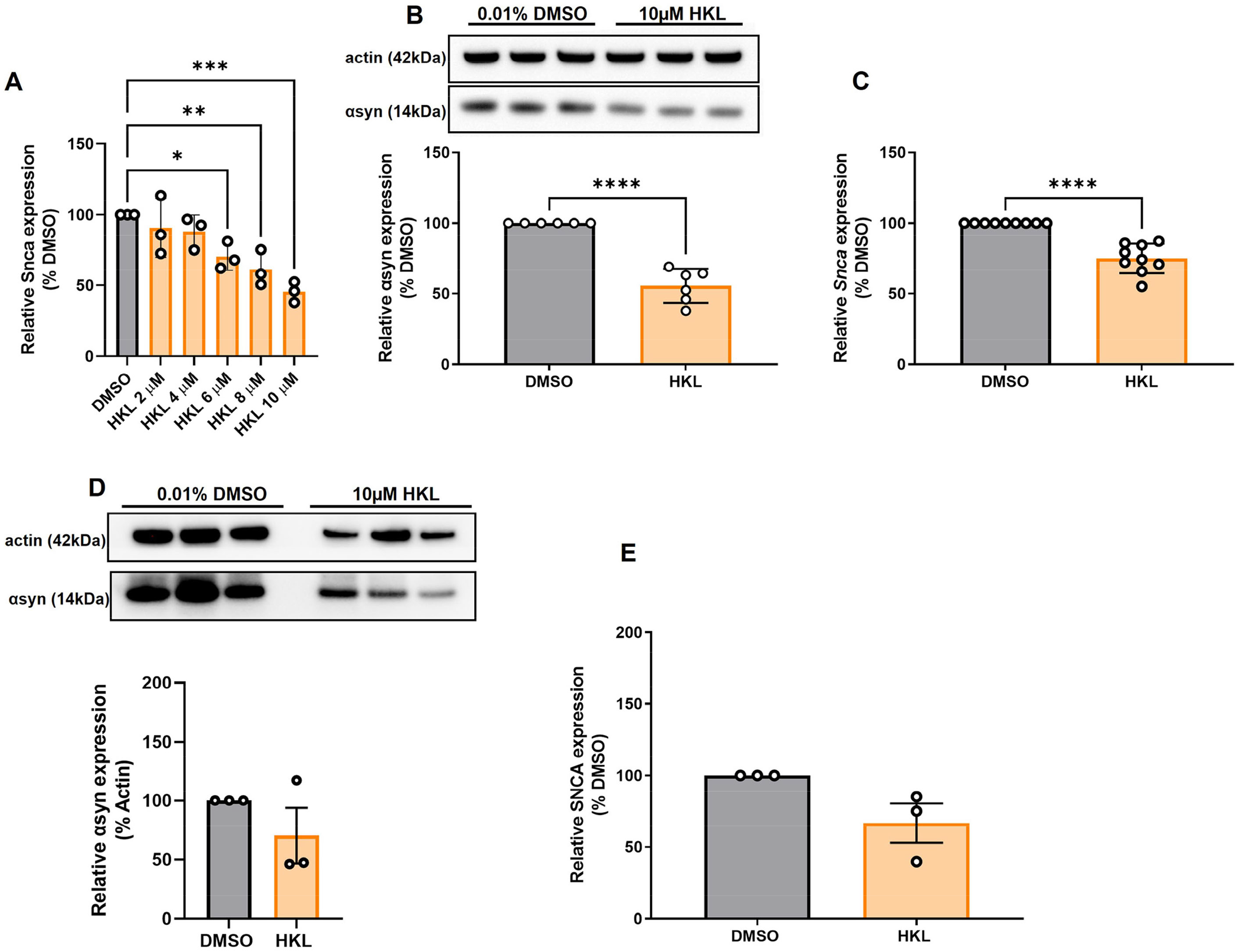
HKL reduces *Snca* mRNA and αsyn levels in mouse primary cortical neurons but not in human-derived iPSC harboring *SNCA* triplication. Verification of effective dose of HKL on *Snca* levels in mouse primary cortical neurons (n = 3 biological replicates/treatment, One-way ANOVA and Dunnet’s post-hoc) (A). Confirmation of αsyn expression reduction (n = 3 biological replicates/treatment) (B) and *Snca* expression reduction (n = 4 biological replicates/treatment) induced by 10µM HKL (C). Western blot and quantification of αsyn expression following 10μM HKL treatment on iPSCs (n = 3 biological replicates/treatment) (D). Levels of *SNCA* mRNA following 10μM HKL treatment (n = 3 biological replicates/treatment) (E). Data are analyzed with unpaired Student’s *t* test and represented as mean ± SEM. *p < 0.05; **p < 0.01, ***p < 0.001.

To confirm the effect of HKL on endogenous human *SNCA* we treated human iPSC-derived neurons harboring the *SNCA* triplication with 10μM HKL for 72 hours. We observed a non-significant 30% reduction in αsyn protein expression induced by HKL (Figure 3D) (*t* (4) = 1.26, p = 0.28) and a non-significant 33.4% reduction in *SNCA* mRNA levels (Figure 3E) (*t* (4) = 2.43, p = 0.07).

### 3.6. Chemical analogues of HKL are not effective in modulating αsyn

Magnolol is a structural isomer of HKL also extracted from the bark of magnolia, differing only by the position of one hydroxyl group (Figure 4A), and has been reported to have similar biological effects as HKL (Hoi, Ho et al. 2010). Hence, we wanted to determine whether magnolol or 4-O-methyl-honokiol (4-0-M-HKL), a HKL-like derivative with good blood brain barrier permeability (Lee, Choi et al. 2011), exhibit similar effects on αsyn regulation. Somewhat surprisingly, we found that magnolol and 4-O-M-HKL have no effect on αsyn (F (3, 8) = 7.79, p < 0.01) and *SNCA* mRNA levels (F (3,8) = 5.91, p < 0.05) (Figure 4B, 4C). This finding highlights the specificity and robustness of HKL in regulating αsyn expression and *SNCA* modulation.

**Figure 4.**
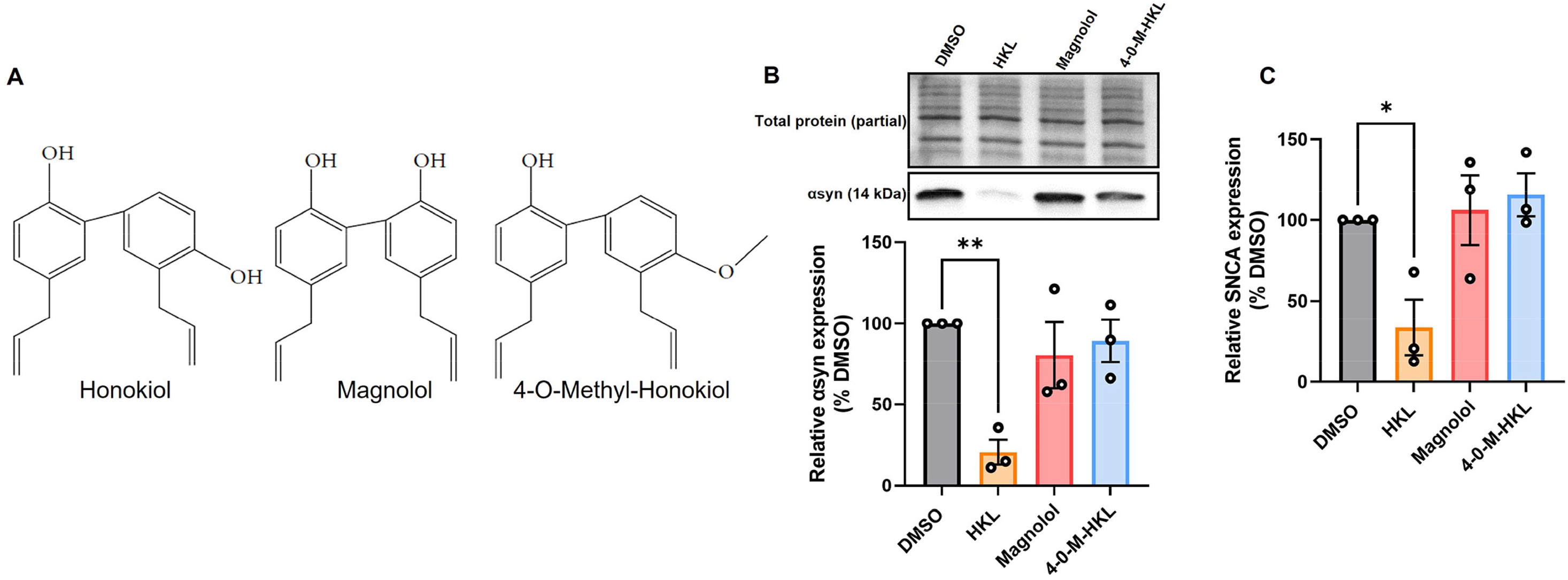
HKL analogues do not modulate expression of αsyn and *SNCA* mRNA in H4 wt-αsyn cells. Structures of HKL, Magnolol, and 4-O-M-HKL (A) Western blot and quantification of αsyn expression following 72h treatment with 10μM HKL, 10 μM Magnolol, and 10 μM 4-0-Methyl-HKL (B). Levels of *SNCA* mRNA following treatment with HKL and analogues (C). Data are analyzed with one-way ANOVA followed by Dunnet’s post-hoc test and are represented as mean ± SEM (n = 3 biological replicates/treatment). *p < 0.05, **p < 0.01.

### 3.7. HKL differentially regulates gene expression

To further assess specific genetic regulatory targets of HKL we conducted bulk RNA sequencing of mouse primary cortical neurons treated with 10µM HKL or vehicle (DMSO) for 72 hours. As expected, transcriptomic analysis confirmed *Snca* down-regulation in HKL treated cells and identified numerous differentially expressed gene (DEG) targets between groups (Figure 5). Combining a discovery and replication dataset from 2 mouse litters revealed a total of 293 DEGs. Importantly, among the top 25 DEGs, three major classes of targets were identified, and these encode for proteins involved in myelination (Bcas1), synaptic transmission and cellular communication (*Angptl4*, *Pla2g7*), and cell signaling and transmembrane transport (*Neat1, Gfap*). To further validate these findings, we selected 4 DEGs and validated the effects of HKL on their expression using quantitative RT-PCR. Consistent with our bulk RNA sequencing data *Angptl4* and *Neat1* were significantly upregulated by HKL and *Snca, Cav1*, and *Kcnq3* were all significantly downregulated by HKL (Supplementary Figure S3). Further investigation and pathway analysis will be required to clarify the specific cellular pathways modulated by HKL that may directly or indirectly lead to the reduction of αsyn levels. These data support HKL as a potential new tool to identify pathways contributing to αsyn pathology and identify potential therapeutic targets.

**Figure 5.**
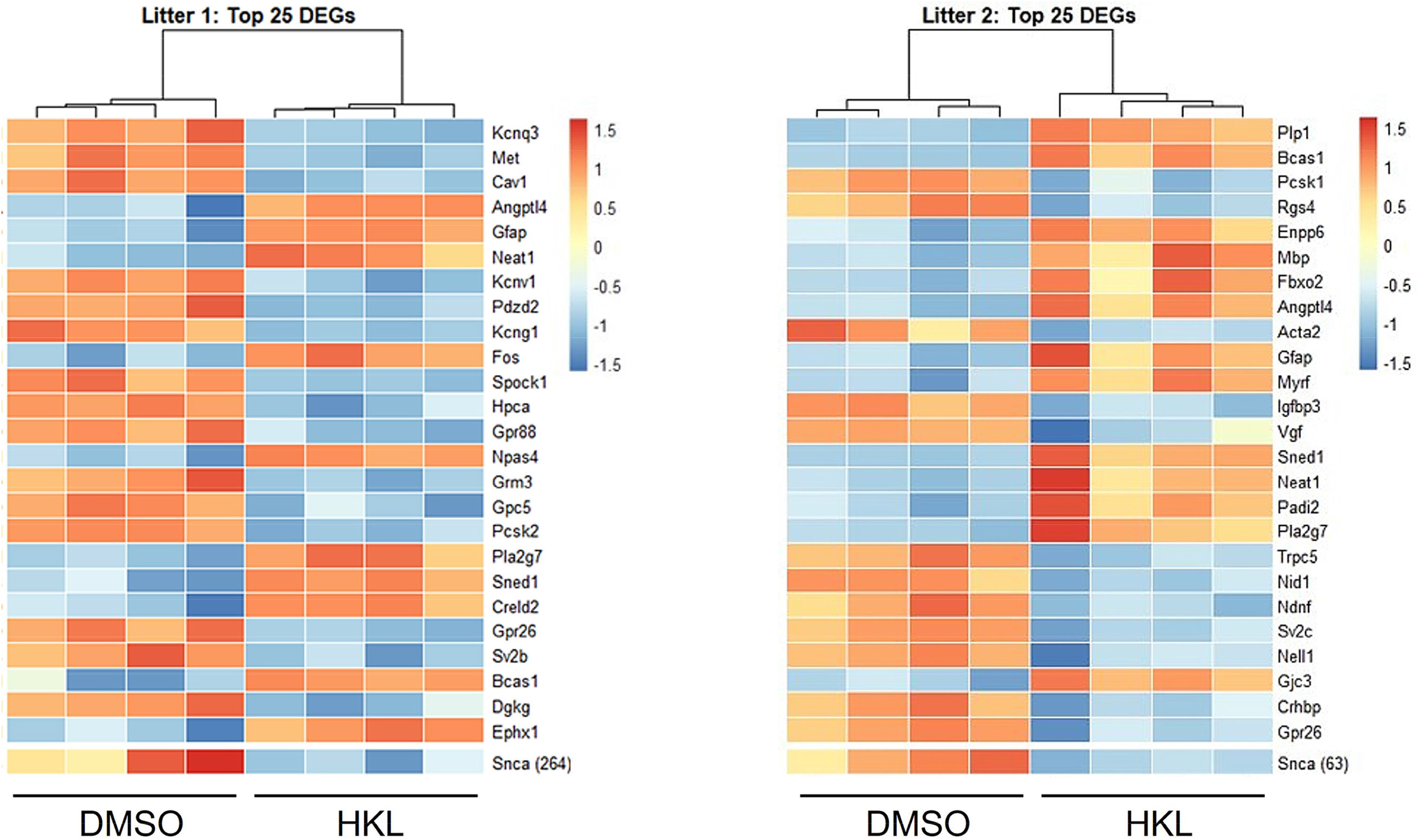
HKL modulates genes associated with myelination, cell signaling and synaptic transmission. Bulk RNA sequencing analysis of 2 litters was performed and revealed 293 differentially expressed genes (DEGs).

### 3.8. HKL reduces rotenone-induced αsyn expression

Because HKL reduces *SNCA/Snca* and αsyn levels in multiple cellular models, we next asked if HKL can modulate αsyn levels under pathological conditions. Abundant data supports the fact that exposure to rotenone, a worldwide-used pesticide, is associated with human parkinsonism and in vitro treatment with rotenone has been widely used to model synucleinopathy while in vivo rotenone treatment can induce a parkinsonian-like phenotype with nigrostriatal degeneration (Sanders et al., 2013; De Miranda et al., 2018). When H4 wt-syn cells are treated with rotenone we observe a significant increase in αsyn protein levels (F (2,18) = 21.08, p < 0.0001) and *SNCA* mRNA expression (F (2,6) = 21.57, p < 0.01) (Figures 6A and 6C). Subsequent treatment with 10µM HKL after rotenone exposure reduced both αsyn protein and *SNCA* expression levels indicating that HKL can prevent a rotenone-induced cellular response (αsyn: *t* (12) = 4.02, p < 0.01; SNCA: *t* (4) = 12.90, p < 0.001) (Figures 6B and 6D).

**Figure 6.**
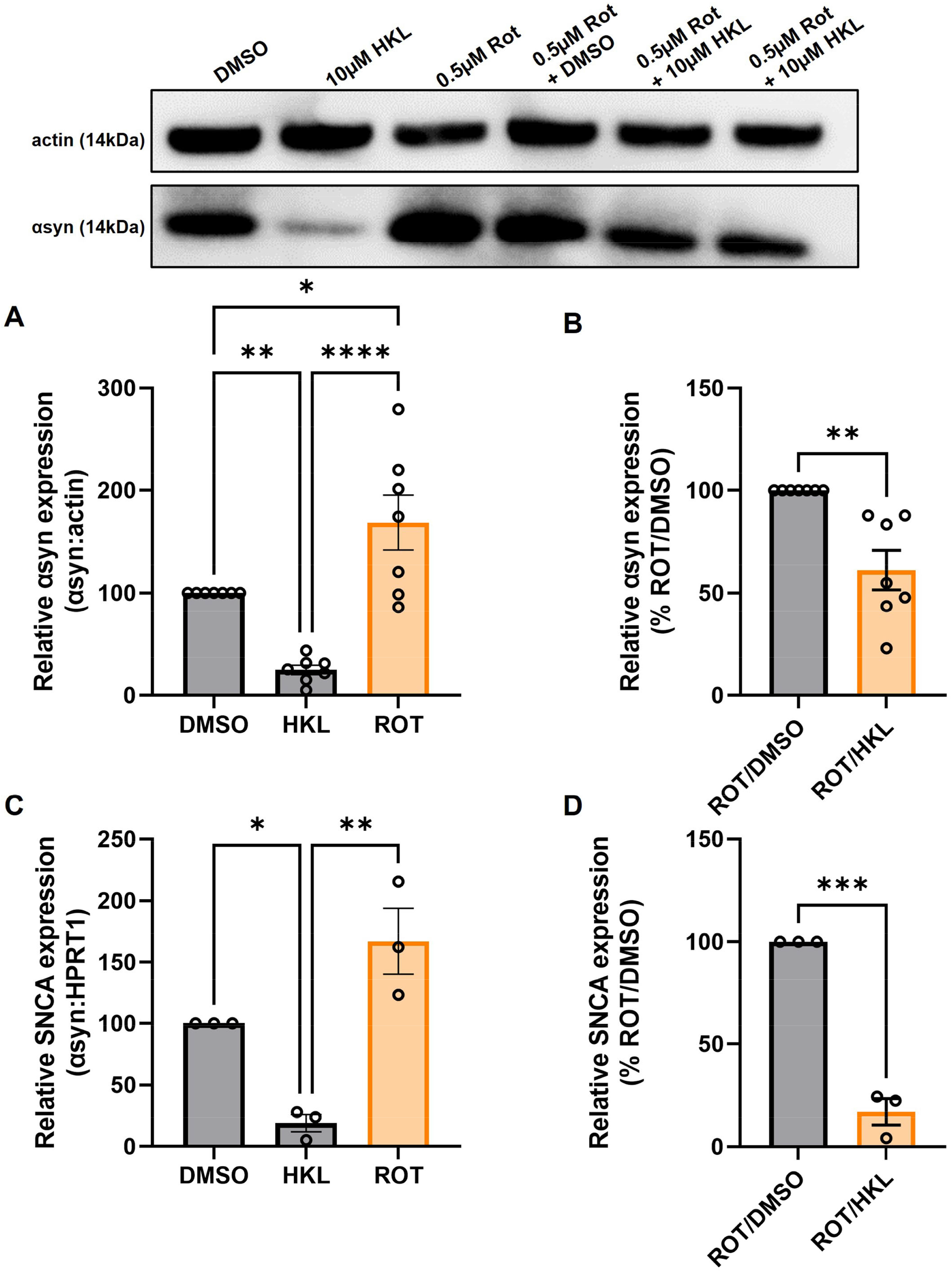
HKL reduces αsyn levels in H4 wt-syn cells treated with rotenone. Western blot and quantification of αsyn expression following treatment with 10µM HKL and 0.5µM rotenone (n = 7 biological replicates/treatment, One-way ANOVA and Dunnet’s post-hoc) (A). Western blot and quantification of rotenone-induced αsyn expression after HKL and DMSO treatment (n = 7 biological replicates/treatment, Student’s *t* test) (B). qPCR assay to determine *SNCA* mRNA levels after 10µM HKL and 0.5µM rotenone treatment (n = 3 biological replicates/treatment, One-way ANOVA and Dunnet’s post-hoc) (C). qPCR analysis of rotenone-induced *SNCA* mRNA levels after HKL and DMSO treatment (n = 3 biological replicates/treatment, Student’s *t* test) (D). Data are represented as mean ± SEM. *p < 0.05; **p < 0.01; ***p < 0.001; ****p < 0.0001.

## 4. Discussion

PD is neuropathologically characterized by intracellular inclusions of aggregated αsyn. Multiplications of the *SNCA* gene locus increases the risk of PD, making αsyn attenuation an important target for drug discovery (Singleton, Farrer et al. 2003; Olanow and Kordower 2017). Current therapeutics for PD include medications to promote dopamine production, such as Levodopa, and surgical interventions such as deep brain stimulation (DBS), for advanced stage patients (Salat and Tolosa 2013; Groiss, Wojtecki et al. 2009). These treatments are useful in reducing the symptoms of the disease but are not effective to slow the progression of the disease.

In the present study, we demonstrate that HKL, a natural, brain permeable small molecule, can successfully reduce αsyn expression in multiple *in vitro* models of PD, which may present a way to slow disease progression. In the human H4 neuroglioma cell line stably overexpressing wt-αsyn under tetracycline regulation, we provide evidence that 10µM HKL treatment is non-toxic and able to efficiently reduce αsyn and *SNCA* mRNA levels (Figures 1A, B, 2A-D, S2A-C). Similar effects are observed in mouse primary cortical neurons (Figure 3A-C) and in patient derived *SNCA* iPSC-derived neurons carrying the PD-associated *SNCA* triplication (Figure 3D, E), although we note that αsyn protein and mRNA level decreases did not quite reach statistical significance in the latter. Our data suggest that the effect of HKL is not mediated by increased rates of protein or mRNA degradation (Figures 1C and 2F). Additionally, we found that the effects on αsyn and *SNCA* mRNA expression are specific to HKL, as they were not reproduced with structurally-related HKL analogues, including Magnolol, an isomer of HKL (Figure 4). Further, we evaluated the effect of HKL in a pathological environment of αsyn overexpression and showed that HKL reverses rotenone-induced overexpression of αsyn and *SNCA* mRNA levels (Figure 6). Finally, we took steps towards identifying a mechanism by which HKL may produce these effects by highlighting genes that are differentially regulated in response to HKL treatment (Figures 5, S3).

Previous research has shown that HKL also prevents formation of αsyn aggregates, possibly by stabilizing αsyn native conformation (Das, Pukala et al. 2018). Additionally, mice with a unilateral 6-OHDA striatal lesion that undergo sub-chronic treatment with HKL demonstrate improvements in motor function, attenuation of nigrostriatal dopaminergic neuronal loss, and reduction in oxidative stress (Chen, Chang et al. 2018). We used H4 cells stably overexpressing αsyn as an *in vitro* synucleinopathy model (Delenclos, Burgess et al. 2019) to triage any effects of HKL on αsyn and *SNCA* mRNA levels. In line with previous results, we observed significant and consistent reduction in αsyn levels (Figures 1B, 2D, 4B, 6A, B) and demonstrated, for the first time, that HKL reduces *SNCA* gene expression (Figures 2A-C, 4C, 6C, D). Notably, in contrast to previous reports of cell cycle arrest and apoptosis induced by HKL in H4 cells after 48-hour treatment (Guo, Ma et al. 2015), we did not observe significant toxicity or changes in proliferation rate associated with HKL treatment (Figure S2A-C).

Importantly, our primary cortical neurons physiologically expressing αsyn under the mouse *Snca* promoter provide a more physiologically relevant neuronal environment and allow us to recapitulate the effects of HKL in reducing αsyn protein and mRNA levels (Figure 3A-C). Finally, using iPSC-derived neurons from patients harboring the *SNCA* gene triplication, we tested the effects of HKL on a physiologically relevant PD model. Our results indicate a non-significant decrease in *SNCA* mRNA and protein levels (Figure 3D, E), most likely because of the small number of biological replicates included in the current study. Additional studies will need to be conducted to clarify the different transcriptional and post-transcriptional effects on the *Snca* and *SNCA* genes. Nonetheless, it is important to note that HKL promotes an overall reduction in αsyn and mRNA levels in different cell models, but the effect size may differ among the models. Therefore, caution is necessary when investigating the mechanisms of in vitro αsyn modulation induced by HKL.

HKL, magnolol, and 4-O-M-HKL are polyphenols with known antioxidant, anti-inflammatory, and anti-tumor effects (Shen, Man et al. 2010), (Liu, Zang et al. 2008), (Lee, Yuk et al. 2009). Recent evidence even proposes these compounds to have neuroprotective potential (Kumar and Khanum 2012). Although these compounds are structurally similar and share mechanisms to exert their effects (Woodbury, Yu et al. 2013), magnolol and 4-O-M-HKL did not modulate levels of αsyn and *SNCA* mRNA in our experiments (Figure 4). This finding suggests that HKL has a unique mechanism of action to regulate αsyn.

It has been suggested that HKL modulates the amyloidogenic pathway by activating Sirtuin-3 (SIRT3) exerting antioxidant activity and improving mitochondrial function (Ramesh, Govindarajulu et al. 2018). The mechanism of HKL in regulating αsyn aggregation has not been clearly elucidated, but recent evidence suggests HKL inhibits fibril formation by directly interacting with lysine-rich region of the N-terminus of the A53T αsyn (Das, Pukala et al. 2018, Jovcevski, Das et al. 2020). Here, we demonstrate that regulation of αsyn and mRNA levels by HKL do not result from increased rates of transcript and protein degradation (Figures 1C 2F), and are probably not a direct regulation of transcription, indicating that HKL could be acting to post-transcriptionally modulate *SNCA* and *Snca* genes, thus expression of αsyn would be reduced.

Our RNAseq results are in line with this hypothesis by demonstrating that the *SNCA* gene was downregulated in cultures from two mouse litters subjected to sequencing (Figure 5). Furthermore, we identified other genes that were differentially expressed after HKL treatment. Of interest, *Angptl4* is an up-regulated gene that encodes angiopoietin-like 4, a secreted protein that modulates triacylglycerol homeostasis (Koliwad, Gray et al. 2012). Indeed, HKL is a partial agonist of peroxisome proliferator-activated receptor-gamma (PPARγ) and was shown to have its neuroprotective effects inhibited by PPARγ antagonists in a hemi-parkinsonian mouse model (Chen, Chang et al. 2018). Considering that PPARγ signaling may influence expression and activity of several genes associated with redox balance, fatty acid oxidation, immune response, and mitochondrial function (Corona, de Souza et al. 2014), it is possible that HKL indirectly modulates αsyn expression via PPARγ activation. Further studies will be required to confirm or refute this hypothesis. Another potential pathway for αsyn modulation induced by HKL is via the long-noncoding RNA *Neat1*, an essential structural component of nuclear paraspeckles that has been found increased in the brains and leukocytes of PD patients (Boros, Maszlag-Török et al. 2020). It is possible that *Neat1* suppresses the expression of hyper edited *Snca* transcripts through nuclear retention, and/or inhibition of nuclear-cytoplasm transport (Prasanth, Prasanth et al. 2005, Chen and Carmichael 2009). Of note, it has been reported that HKL is a modulator of sirtuin3 and other AMPKα-CREB signaling pathways (Ramesh, Govindarajulu et al. 2018). However, we have not observed significant changes related to Sirtuin3 and AMPKα-CREB signaling-associated genes in our cellular models following HKL treatment (unpublished data).

In summary, we demonstrate that the natural small molecule HKL can modulate levels of αsyn protein and transcript in multiple cell models of synucleinopathies. We also provide initial evidence of mechanisms by which HKL regulates αsyn, which suggests both direct gene regulation and indirect metabolism regulation. Additional studies will need to validate these findings in *in vivo* PD models and to clarify pathways through which HKL reduces αsyn.

## 6. Author Contributions

MD and PJM contributed to the conception and design of the study. SF, JDB, ML, DA, ZK, AHF, IS, DJQ PP, and NND conducted experiments, collected, and analyzed the data. SF wrote the first draft of the manuscript. SLB contributed to data analysis and the first draft of the manuscript. SF, SLB, PJM, JDB, and NND contributed to manuscript revision. All authors contributed to manuscript revision, read, and approved the submitted version.

## 8 Acknowledgements

We would like to thank Christine Perez-Rosa and Morgan Russ for their technical support on this project as well as the Mayo Clinic Genome Analysis Core for performing the RNA sequencing and analysis.

---

**Supplemental Figure S1.**
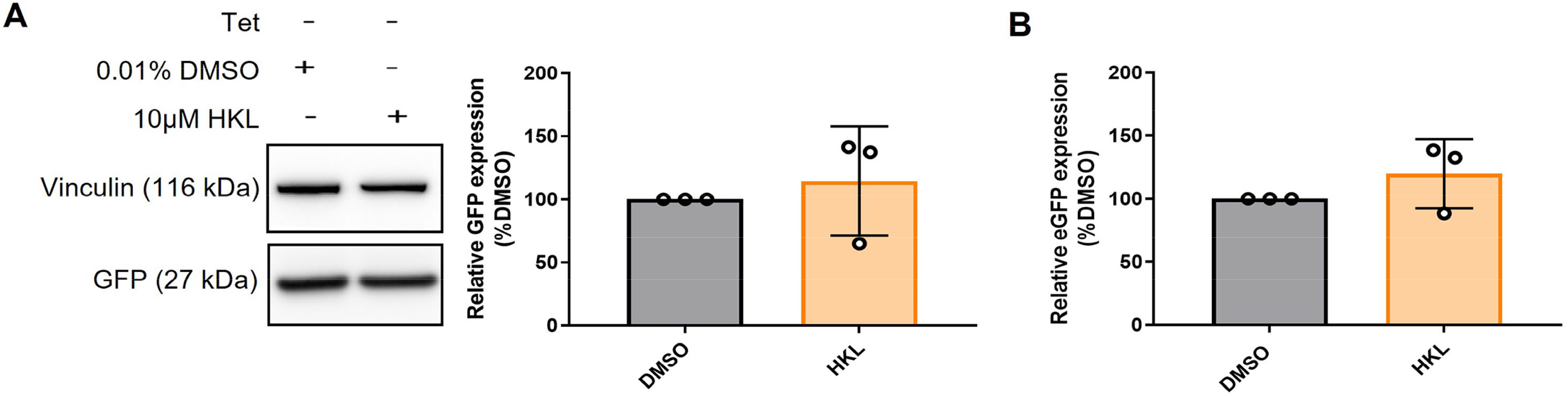
HKL does not affect GFP expression in H4 cells stably overexpressing wt-αsyn. Western blot and quantification of GFP expression following treatment with 10µM HKL (n = 3 biological replicates/treatment, *t* (4)=0.58, p = 0.59) (A). Effects of HKL treatment in eGFP mRNA levels (n = 3 biological replicates/treatment, *t* (4)1.25, p = 0.28) (B). Data are analyzed with unpaired Student’s *t* test and are represented as mean ± SEM.

**Supplemental Figure S2.**
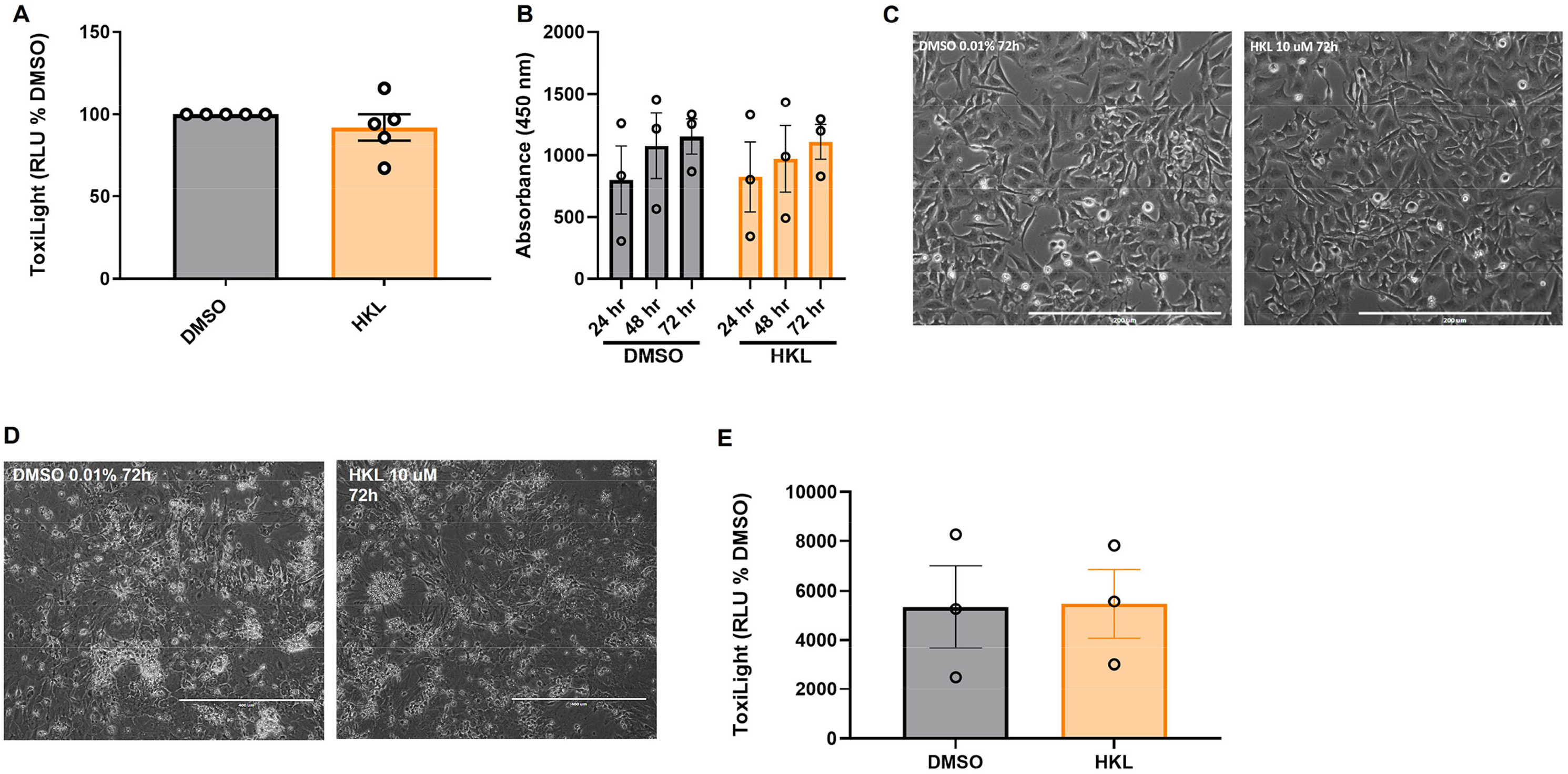
HKL does not induce toxicity nor affect cell proliferation. The Toxilight assay was performed in H4 wt-αsyn cells after 72h treatment with 10µM HKL (n = 5 biological replicates, *t* (8) = 0.10, p = 0.35) (A). Proliferation WST-1 assay in H4 wt-αsyn cells assessed over 72h of HKL treatment (n = 3 biological replicates, repeated measures two-way ANOVA, time F (1.13, 9.03) p < 0.05, treatment F (3, 8) = 0.35 p = 0.35, time X treatment F (6, 16) = 0.17 p = 0.98) (B). Toxicity of 72h treatment with 10µM HKL on mouse primary cortical neuron was evaluated in the Toxilight assay (n = 3 biological replicates, *t* (4) = 0.06, p = 0.96) (C). Data are analyzed with unpaired Student’s *t* test and are represented as mean ± SEM.

**Supplemental Figure S3.**
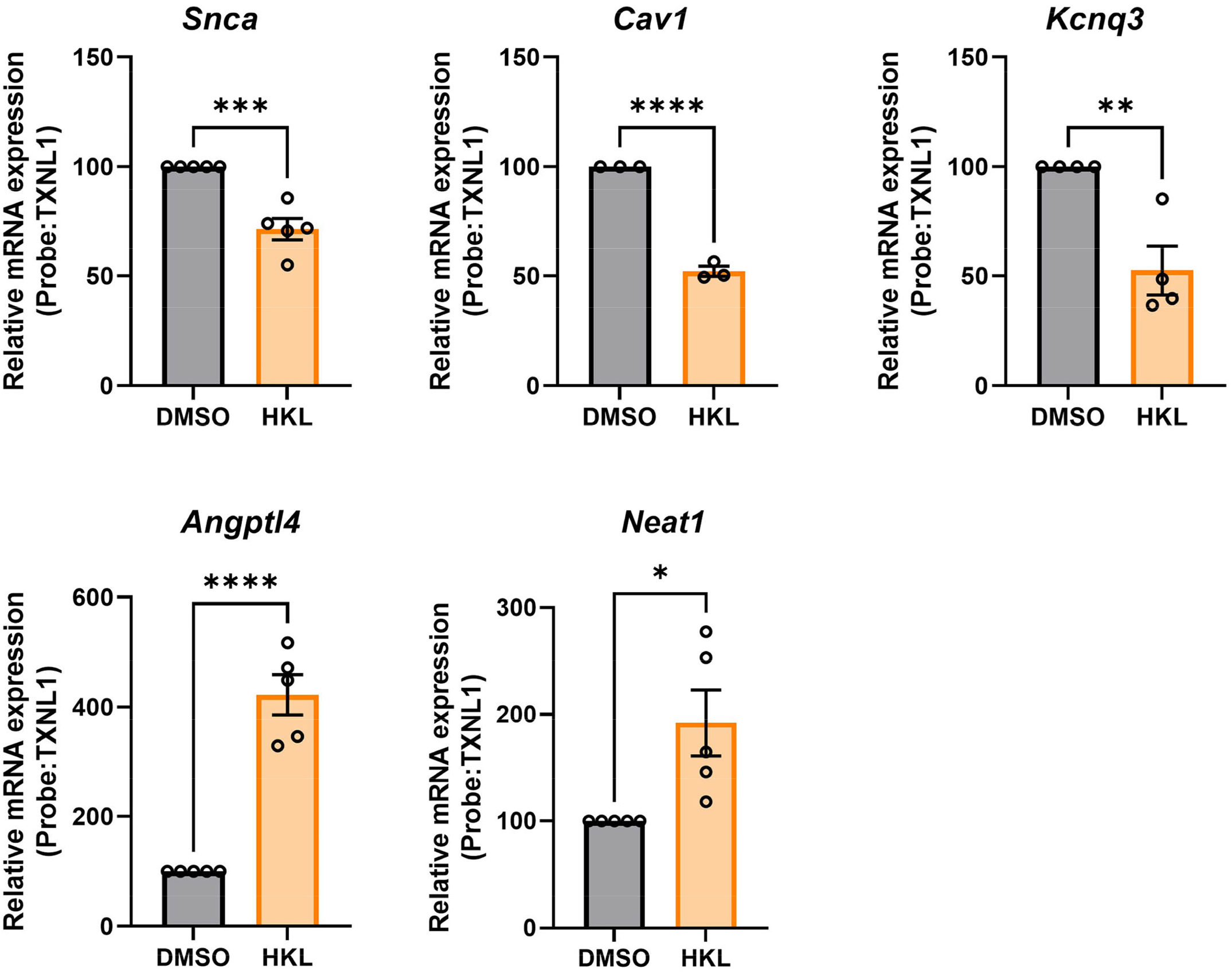
Validation of DEGs in mouse primary cortical neurons. Of the top 25 differentially expressed genes (DEGs) identified with bulk RNA sequencing in mouse primary cortical neurons, the following were validated to confirm the effect of HKL in these cells. Data were analyzed with Student’s t test and are represented as mean ± S.E.M. ****Cav1 -*t* (4) = 21.12, p = <0.0001, **Kcnq3 - *t* (6) = 4.21, p = 0.0056, ****Angplt4 - *t* (8) = 8.82, p = < 0.001, *Neat1, *t* (8) = 2.95, p = 0.0184, ***Snca *t* (8) = 5.79, p = 0.0004.

